# Stereotypy is strongly linked to multiple biomarkers of oxidative stress—a potential common etiology for Abnormal Repetitive Behaviors

**DOI:** 10.1101/2025.06.09.658735

**Authors:** Kendall M. Coden, Kaleigh J. Beacham, Beatriz E. Stix-Brunell, Roberta Moorhead, Kyna A. Byrd, Joanna N. Baker, Jerome T. Geronimo, Karen J. Parker, Joseph P. Garner

## Abstract

Spontaneous stereotypies (abnormal, repetitive, and seemingly goal-less behaviors) in captive animals resemble stereotypies documented in patients with neurodevelopmental disorders, including evidence of homologous cortico-striatal dysfunction and shared behavioral deficits. While environmental risk factors for stereotypies are well documented, their developmental pathophysiology remains unclear. However, as previously found for compulsive behavior, there is growing evidence that REDOX imbalance may be linked to stereotypy. To examine the nature of this relationship, we first tested whether plasma glutathione level, the gold-standard biomarker of REDOX imbalance, is predictive of stereotypy severity in N=19 C57BL/6 mice. After confirming the presence of this relationship, we used a proteomics approach (Olink) to identify a broader biomarker profile of dysfunction. We found expression of 9 proteins to correlate with plasma glutathione level, and expression of 15 proteins to correlate with stereotypy severity. A subset of these proteins additionally correlated with stereotypy severity in a validation cohort of CD1 mice (N=28). Further supporting a role for REDOX imbalance in the developmental pathophysiology of stereotypies, the identified proteins were associated with REDOX physiology, dopamine physiology, and stereotypy-presenting human neurodevelopmental disorders. These data suggest REDOX imbalance may contribute to the developmental pathophysiology of abnormal repetitive behaviors and highlight promising novel targets for intervention.

## Introduction

Stereotypies (abnormal, repetitive, unvarying, seemingly goal-less behaviors) are prevalent within the animal kingdom and have been documented in nearly every captive mammal and bird species, including laboratory animals (1,2), zoo animals (3,4), and farm animals (5). Additionally, stereotypies are a core feature of several neurodevelopmental and neuropsychiatric disorders in humans such as autism spectrum disorder (ASD; (6,7)) and schizophrenia (1,8)). We have previously established that spontaneous stereotypies in laboratory animals serve as a model of stereotypies observed in humans, with homologous basal ganglia cortico-striatal dysfunction (9–12) and shared behavioral deficits across species (e.g., recurrent perseveration (13) and high levels of knowledge-action dissociation (1)). Despite well documented environmental risk factors associated with the development of stereotypies in captive animals (14,15), the developmental pathophysiology of these behaviors remains elusive.

We and others have previously hypothesized and shown that the manifestation of specific ARB phenotypes is dependent upon which basal ganglia cortico-striatal loop is impaired (6,11,12). The basal ganglia are a group of midbrain nuclei which regulate initiation, termination, and sequencing of both voluntary and automatic motor behavior (16–18) via several reciprocal projections (i.e., loops) with cortical regions (16,19–21). Each of these loops is comprised of a direct (activating) and indirect (inhibitory) pathway which work in opposition to regulate behavior (16–18), with dopamine modulating the balance between the two (22) . Dysfunction of the indirect pathway is associated with production of abnormal repetitive behaviors (ARBs) including stereotypies and compulsive behaviors (13,23–25). For compulsive behaviors, there is an established causal relationship between REDOX imbalance (a state in which physiological demand for antioxidants [e.g. glutathione (GSH)] surpasses their bioavailability), and development, symptom severity, and treatment of the ARB (26–28). Although the developmental pathophysiology of stereotypy remains unknown, the homologous, highly conserved nature of the basal ganglia loops (6,29), suggests that REDOX imbalance may also contribute to the developmental pathophysiology.

Despite evidence implicating REDOX imbalance in several neurodevelopmental, neurodegenerative, and psychiatric conditions (30–34), to our knowledge, no research has directly assessed the relationship between REDOX imbalance and stereotypy. Therefore, we aimed to test the hypothesis that stereotypies, like other ARBs, are related to REDOX imbalance. To examine this, we first tested whether plasma GSH level, a gold-standard biomarker of REDOX imbalance, predicted stereotypy severity in C57BL/6 mice. After confirming this relationship, we next used a state-of-the-art proteomics approach (Olink Mouse Exploratory Panel) to establish a broader biomarker profile of dysfunction within the same group of C57BL/6 mice. Notably, due to the high evolutionary conservation of the proteins, human antibodies are used to quantify 90 of the 96 targets on the panel. We predicted that expression of proteins associated with REDOX balance would be correlated with plasma GSH level and/or stereotypy severity. To ensure our findings were not strain specific, we opportunistically assessed Olink protein profiles in an independent cohort of CD1 mice with stereotypy scores and banked blood samples from an independent study in our lab.

## Results

To increase diversity in stereotypy severity in our sample population, we selected N = 20 C57Bl/6 mice ranging in age from 140 to 225 days old that were retired from other studies and slated for euthanasia. Animals slated for euthanasia due to health concerns (such as meeting humane endpoint for their initial study) were excluded. This approach allowed us to obtain a more heterogenous population of animals, thus increasing translatability and replicability of results (34,35). Additionally, given the heterogeneity present in neurodevelopmental disorders, selection of non-transgenic animals ensures broader heterogeneity in our sample population, and thus, further generalization of results (35). To determine stereotypy severity, we unobtrusively recorded mouse home cage behavior and quantified behavior using a validated mouse ethogram (36). To quantify plasma GSH level, whole blood was collected *via* cardiac puncture and centrifuged to obtain plasma. A commercially available kit was then used to quantify plasma GSH level. One subject was excluded from analyses due to hemolysis of the sample (Final N = 19).

We first investigated whether variation in plasma GSH level predicts stereotypy severity in C57BL/6 mice. To do so, we used a least squares general linear model (LS-GLM). As is necessary in an experiment with heterogeneity in sample population, we included age of animals in our analysis and found an interaction between age and plasma GSH level (F_1,12_ = 6.4668; P = 0.0258). We next examined the age distribution of animals and noted that the animals divided cleanly into two age groups: young mice (n = 10), ranging in age from 140 to 169 days of age, and old mice (n = 9), ranging in age from 185 to 225 days of age, at time of euthanasia. As such, we performed this analysis again using two age groups and found that plasma GSH level positively predicted severity of stereotypy in the young group of mice (T_12_ = 3.6467; P = 0.0033), but not in the old group of mice (T_12_ = -0.7191; P = 0.4858; fig 1). This result suggests that while plasma GSH level is predictive of stereotypy severity, the predictive nature of this relationship diminishes with age.

**Figure 1:**
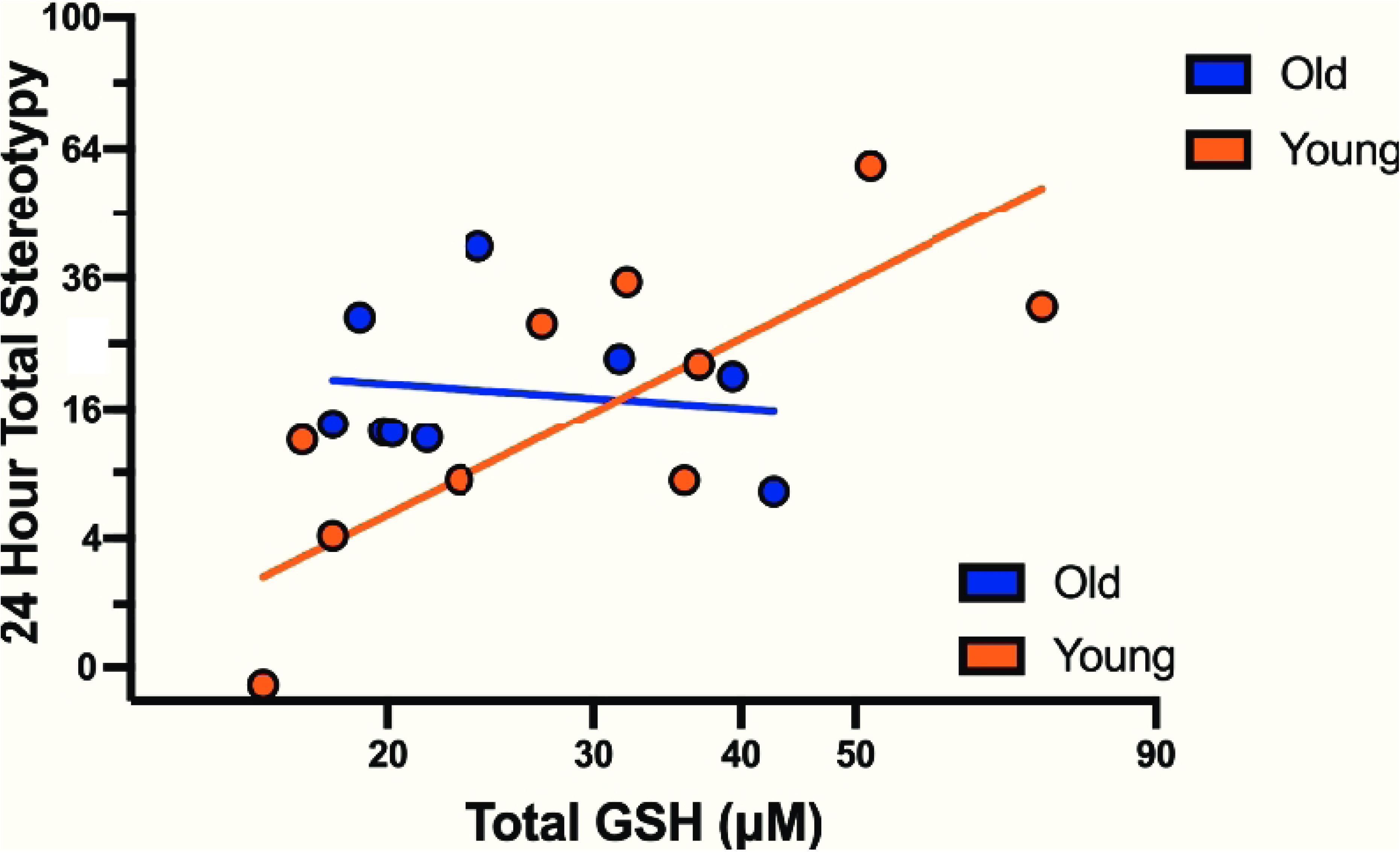
Plasma GSH level positively predicts stereotypy severity in the young group (n = 10; 140-165 days old), but not in the old group (n = 9; 180-225 days old), of C57BL/6 mice. Data are corrected for blocking factors in the analysis. Circles on the graph depict individual mice. Lines on the graph depict regression lines for the two age groups.

After establishing the relationship between our gold-standard biomarker, GSH, and stereotypy severity, we next investigated whether there was a broader biomarker profile that could be used to predict REDOX imbalance and stereotypy severity. To do so, we first used the Olink T96 mouse exploratory panel to characterize the proteomic profile of the C57BL/6 mice. We then examined these proteomic profiles to identify novel plasma protein biomarker profiles associated with plasma GSH level and/or stereotypy severity. Due to low plasma availability, one subject was excluded from analyses (final N = 18). In these analyses, “protein” refers to the average proteomic profile across all subjects, while interactions with protein test how proteomic profile changes with age group, plasma GSH level, and their interaction.

Using two separate restricted maximum likelihood (REML) mixed models controlled for age group, we found a significant correlation between protein expression and plasma GSH level as a function of age (F_91,1274_ = 1.50; *P* = 0.0023); and a significant relationship between protein expression and stereotypy severity (F_91,1274_ = 3.29; *P* < 0.0001). Additionally, there was no evidence of an age group dependent relationship between protein expression and stereotypy severity (F_91,1274_ = 1.01; *P* = 0.4546). Of the 92 proteins analyzed, expression of nine proteins (*Tnni3*, *Dlk1*, *Yes1*, *Ddah1*, *Pdgfb*, *Prdx5*, *Parp1*, *Ghrl*, *and Ahr*) was significantly correlated with plasma GSH level (fig 2A) and expression of 15 proteins (*Tnni3, Ddah1, Riox2, Nadk, Plin1, Gcg, Qdpr, Yes1, Plxna4, Ahr, Parp1, Prdx5, Fst, Ccl20, Pdgfb*) significantly correlated with stereotypy severity (fig 2B). We performed a Benjamini-Hochberg (BH) correction to account for potential false discovery (see fig 2, supplementary table S1 online, and supplementary table S2 online for BH corrected q-values and associated P-values for plasma GSH level and stereotypy severity respectively).

**Figure 2:**
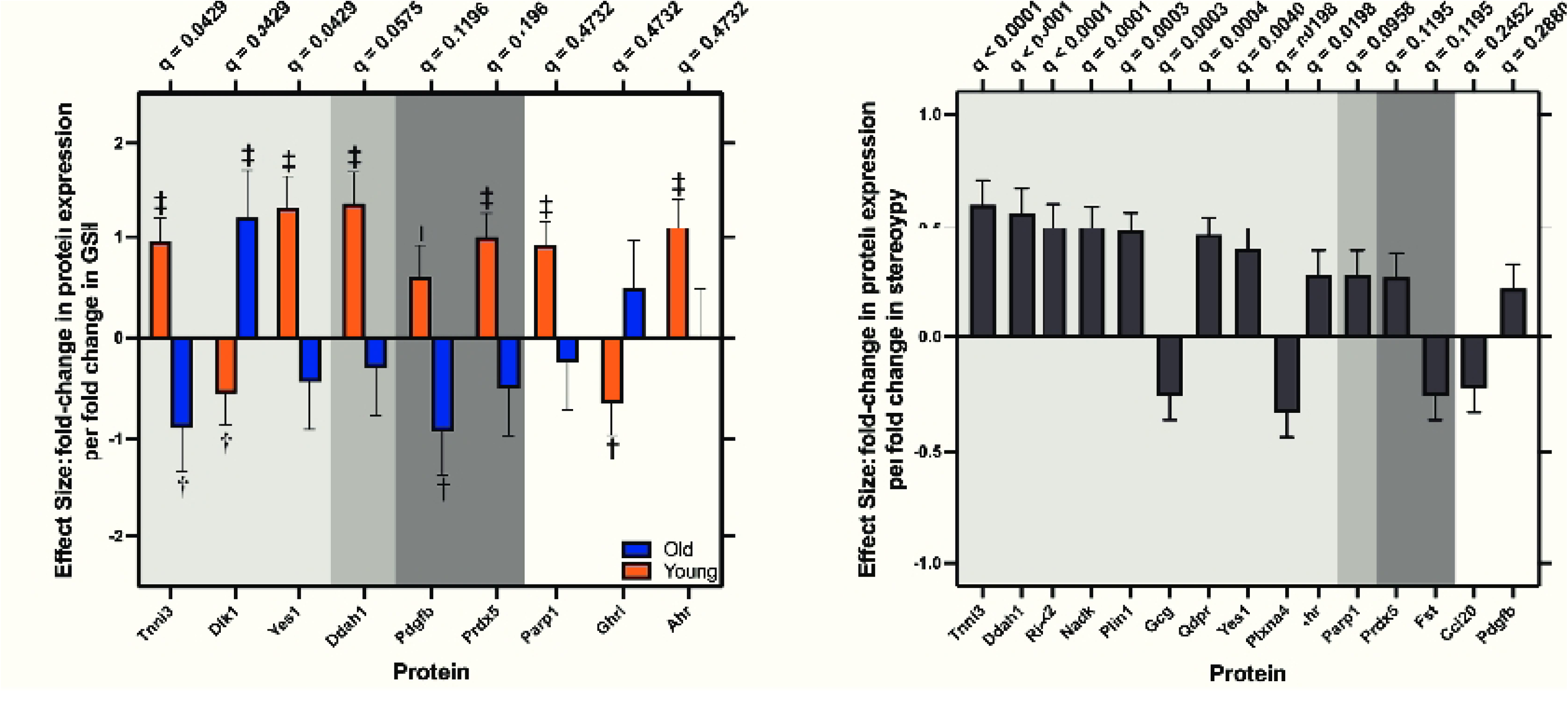
Protein expression predicts both plasma GSH level and stereotypy severity in n = 19 C57BL/6 mice (n = 10 young mice; n = 9 old mice). Regression slope +/-standard error. **a)** expression of nine proteins significantly correlated with GSH level as a function of age. The light gray depicts proteins that pass BH correction at critical q = 5%; medium gray denotes proteins that pass BH correction at critical q = 10%; dark gray depicts proteins that pass BH correction at 20%. Orange bars depict protein expression in the young group of mice and blue bars depict protein expression in the old group of mice. ‡ denotes proteins that pass BH correction at critical q = 5% BH correction within age group; † denotes protein that passes BH correction at critical q = 20% within age group. **b)** Expression of 15 proteins correlated with stereotypy severity once age is taken into account. The light gray depicts proteins that pass BH correction at critical q = 5%; medium gray denotes proteins that pass BH correction at critical q = 10%; dark gray depicts proteins that pass BH correction at 20%.

Given the age-dependent correlation between protein expression and plasma GSH level, for the nine proteins that correlated with plasma GSH level, we separately tested the correlation of individual protein expression and plasma GSH level for young and old groups of animals—the resulting p-values were also BH corrected to q-values. As shown in figure 2A, for the young group of mice, plasma GSH level was positively correlated with expression of *Tnni3*, *Yes1*, *Ddah1*, *Pdgfb*, *Prdx5*, *Parp1*, and *Ahr*, whereas plasma GSH level negatively correlated with expression of *Dlk1* and *Ghrl*. In the old group of mice, expression of *Dlk1* was positively correlated with plasma GSH level, while expression of *Tnni3* and *Pdgfb* were negatively correlated with plasma GSH level (fig 2a; see supplementary table S3 online for age-specific P-values and BH corrected q-values).

To assess the replicability of our proteomic findings and to ensure they were not strain specific, we used the Olink mouse exploratory panel to quantify the proteomic profile of banked plasma samples from 28 CD1 mice from a prior independent study, in which stereotypy severity were also quantified. Using an REML repeated measures mixed model, we found a significant correlation between stereotypy severity (*F_91,2366_* = 1.37; *P* = 0.0133) and protein expression in CD1 mice (fig 3; see supplementary table S4 online for BH corrected p-and q-values), including three proteins (*Pdgfb*, *Riox2*, and *Parp1*), which also correlated with stereotypy severity in C57BL/6 mice. Thus, these results indicate that our proteomic findings are not strain specific. As all of the CD1 mice fell within the age range of young mice, age group analyses were not conducted.

**Figure 3:**
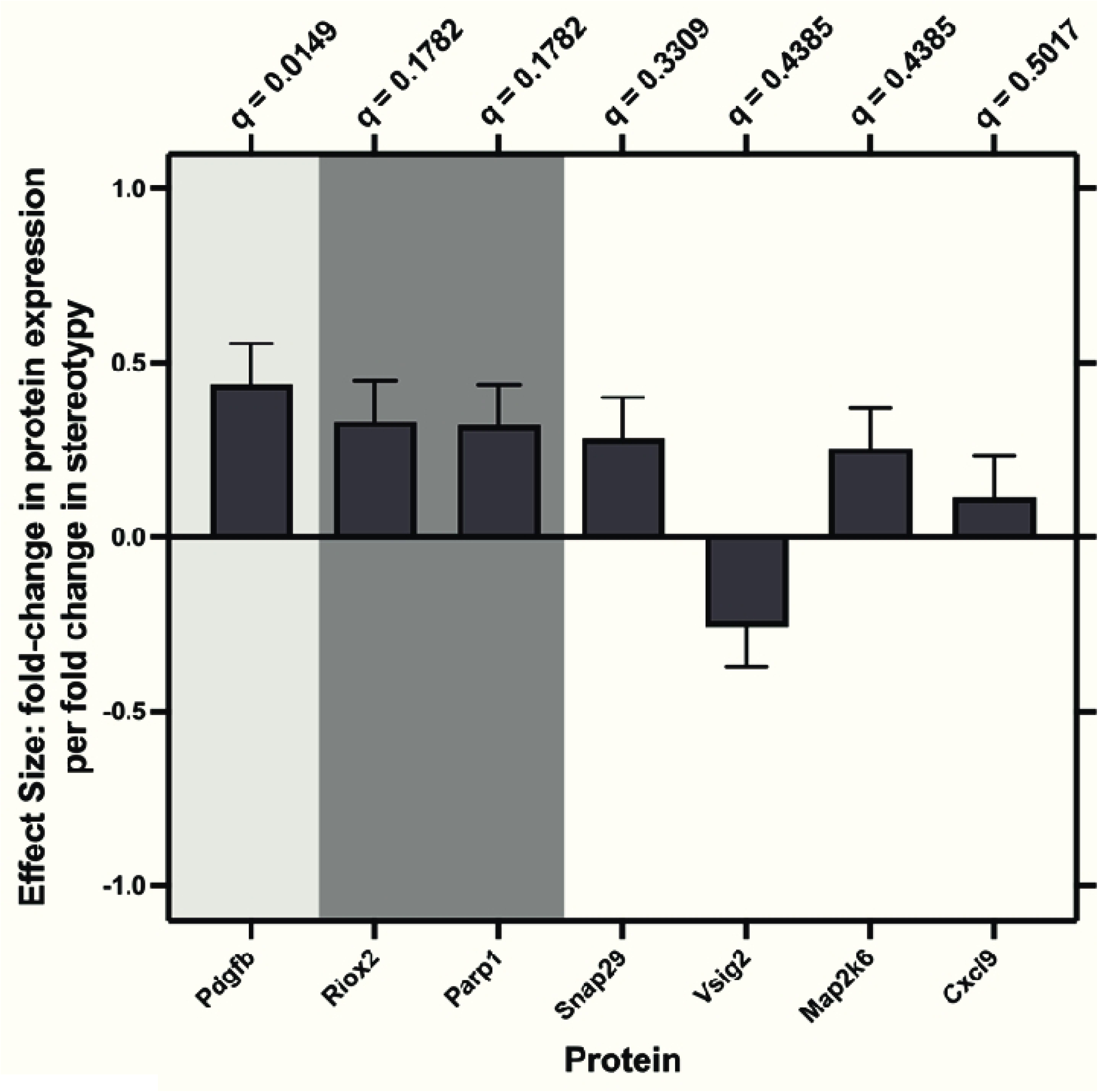
Expression of proteins significantly correlated with stereotypy severity in CD1 Mice (n = 28). Regression slope +/-standard error. Expression of 7 proteins significantly correlated with stereotypy severity in CD1 mice. Light gray shading denotes protein passed BH correction with critical q = 5%; darker gray denotes protein passes BH correction at critical q = 20%. Expression of *Pdgfb*, *Riox2*, *Parp1*, *Snap29*, *Map2k6*, and *Cxcl9* increase with severity of stereotypy, whereas expression of *Vsig2* decreases with stereotypy severity. Expression of *Pdgfb*, *Riox2*, and *Parp1* is also correlated with plasma GSH level and stereotypy severity in C57BL/6 mice, indicating that they are biomarkers of REDOX balance and stereotypy severity.

We next examined the functional roles of identified protein biomarkers which were significant in at least one of these analyses and passed BH correction for q < 20%. Of the 15 proteins that met criteria, three were REDOX physiology related, one was dopamine related, and 11 were related to both dopamine and REDOX (see fig 4a for classifications). Noting the interaction between age group and the predictive nature of plasma GSH level on stereotypy severity, and the relationship between age group, plasma GSH level, and protein expression level, we next examined whether the proteins were known biomarkers of neurodevelopmental and neuropsychiatric disorders. We strategically chose a pediatric onset disorder characterized by stereotypies (ASD), an adolescent/early adulthood onset disorder also characterized by stereotypies (schizophrenia), and a pediatric onset disorder not characterized by stereotypies (attention-deficit hyperactivity disorder—ADHD).

**Figure 4:**
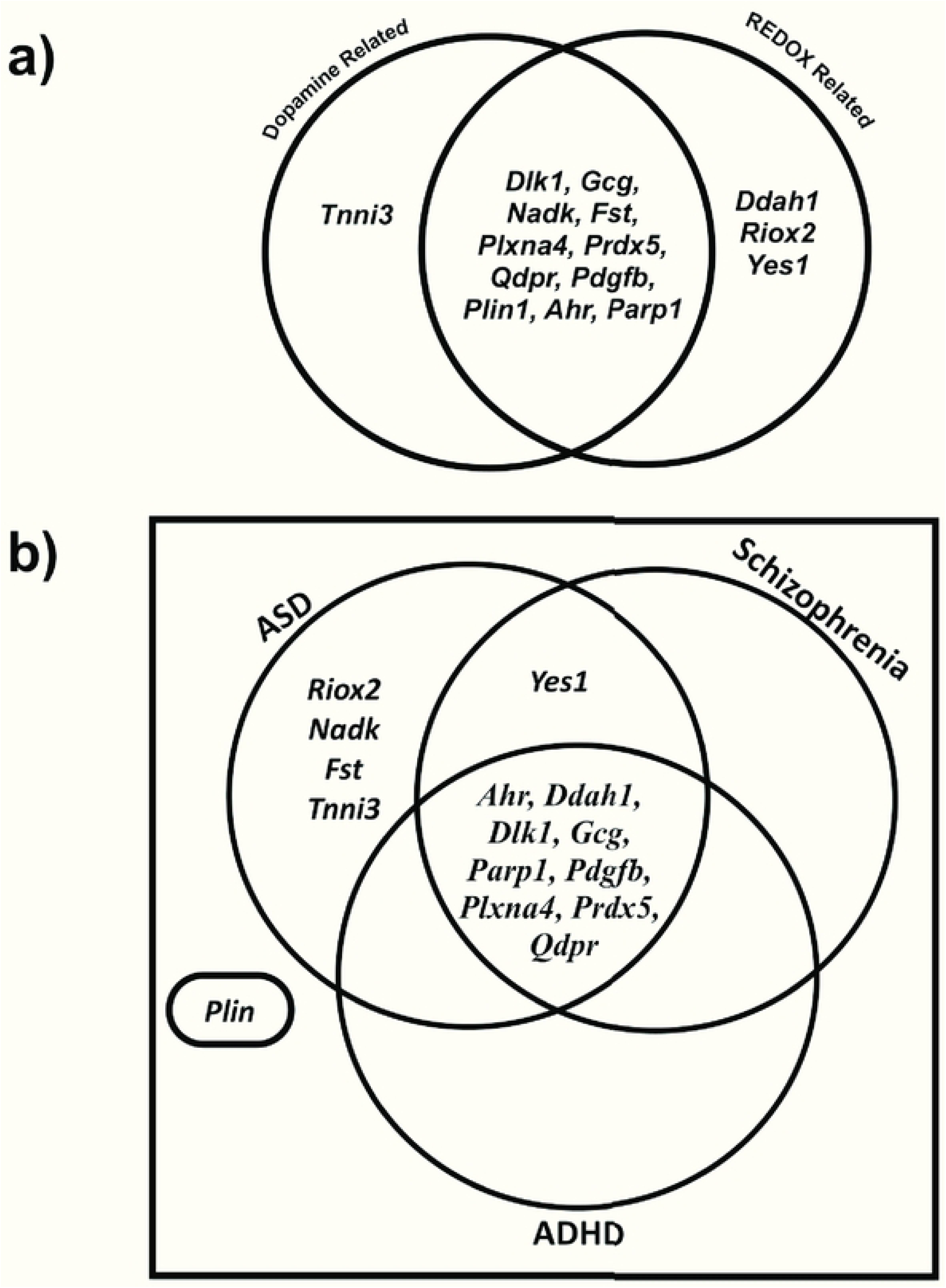
Ontology of identified proteins which passed BH correction of q < 20% in at least one analysis in either the C57BL/6 (n = 18) or CD1 mice (N = 28). a) Venn diagram depicting proteomic “hit” associations with REDOX physiology (far right section), dopamine physiology (far right section), or both REDOX and dopamine physiology (middle section). b) Venn diagram depicting association of proteomic “hits” as biomarkers for three disorders (ASD, Schizophrenia, and ADHD). Proteins associated with multiple disorders are located in overlapping region(s) enclosed by disorder specific circles. Proteins located outside of the Venn diagram are not associated with any of these disorders.

We found that the number of proteins associated with each respective disorder changed in response to both age of disorder onset as well as absence or presence of stereotypies (Fig 4b). For ASD, a pediatric onset disorder characterized by presence of stereotypies, 14 of the 15 identified proteins (i.e., *Riox2, Nadk, Fst, Tnni3, Yes1, Ahr, Ddah1, Dlk1, Gcg, Parp1, Pdgfb, Plxna4, Prdx5, Qdpr;* see fig 4b) are known biomarkers and/or currently under investigation as potential disorder related biomarkers. As for schizophrenia, an adolescent or early adulthood onset disorder characterized by presence of stereotypies, 10 of the 15 identified proteins (i.e. *Yes1, Prdx5, Ahr, Ddah1, Dlk1, Gcg, Parp1, Pdgfb, Plxna4, Qdpr; S*ee fig 4b) are disorder associated biomarkers. As for ADHD, a pediatric onset disorder typically lacking stereotypies but characterized by presence of other forms of ARB, nine of the 15 identified proteins are disorder associated biomarkers (i.e., *Prdx5, Ahr, Ddah1, Dlk1, Gcg, Parp1, Pdgfb, Plxna4, Qdpr*; see fig 4b).

## Discussion

Despite the pervasive nature and extensive literature on stereotypies, the underlying developmental pathophysiology remains poorly understood. Here, we investigated whether REDOX imbalance, a known contributing factor to other forms of ARBs regulated by adjacent basal ganglia circuitry, is also associated with stereotypy severity. Given the growing literature implicating REDOX imbalance in other ARBs (26,27,31) we predicted that plasma GSH level would positively predict stereotypy severity in mice.

After establishing the relationship between REDOX imbalance and stereotypy severity, we used a state-of-the-art proteomics approach (Olink T96 Mouse Exploratory Panel) to identify broader biomarker profiles of REDOX imbalance and stereotypy severity. As predicted, we found strong relationships between protein expression (specifically in proteins related to REDOX imbalance) and both GSH level and stereotypy severity in C57BL/6 mice. We additionally replicated these proteomics findings in a separate cohort of CD1 mice, indicating that these profiles are replicable and robust, as well as independent of strain. Taken together these data indicate that REDOX imbalance may contribute to the developmental pathophysiology of stereotypy.

Mirroring prior evidence implicating REDOX imbalance in the developmental pathophysiology of compulsive behaviors (26,27), in C57BL/6 mice we found a strong positive correlation between plasma GSH level and stereotypy severity in the young, but not in the old, group of mice. There are two potential explanations for the lack of predictive nature of plasma GSH level on the stereotypy severity in old mice: 1) the old mice are no longer experiencing REDOX imbalance, or 2) the lack of signal in GSH is due to other factors producing high levels of stereotypies despite low plasma GSH level. If the first explanation is correct, then we would expect less severe stereotypies in old mice with low plasma GSH levels. However, this result was not found. Rather, when examining proteomic profiles of the C57BL/6 mice, we found one set of proteins to correlate with plasma GSH level in an age dependent manner—the majority (six out of nine) being significantly correlated with plasma GSH level in the young, but not in the old, group—and another set to correlate with stereotypy severity, independent of age. Together, these results favor the second explanation: other factors are contributing to stereotypies in old mice— likely in an additive effect to REDOX imbalance. Clinical evidence further supports this second explanation. A 2012 study (37) found that N-acetylcysteine, a dietary precursor to GSH, reduced repetitive behaviors in younger children with autism (average age 7.0 years +/-2.0 years), while a 2024 study (38) in slightly older children (ranging in age from 8 to 12 years old) found no correlation between central GSH levels and stereotypy, consistent with age-dependent differences in underlying mechanisms.

Further implicating REDOX imbalance in the developmental pathophysiology of stereotypies, proteomic ontology revealed that identified proteomic biomarkers largely correlated with REDOX physiology, dopamine physiology, or both REDOX and dopamine physiology. This finding was largely unsurprising as metabolism of dopamine produces large quantities of neurotoxic radical oxygen species which can be neutralized by antioxidants (22), and dopamine-induced apoptosis is prevented by GSH in a dose-dependent manner (39). Additionally, the identified biomarkers were tightly related to disorders characterized by presence of stereotypies with an influence of age. There is a long-standing hypothesis that stereotypies are behavioral scars resulting from an unknown cause (15). Our findings support this hypothesis and suggest REDOX imbalance as a contributing factor. Taken together, our results suggest that plasma GSH level and/or expression levels of our identified protein biomarkers may serve as risk predictors for developing stereotypies and other ARBs.

While the results presented in this paper are correlational in nature, given the highly conserved cytoarchitecture of basal ganglia loops (16,21) and the causal relationship between REDOX imbalance and compulsive behaviors (26–28) it is likely that REDOX imbalance is also causally related to the development of stereotypy. Although it warrants further investigation, we propose REDOX imbalance as a shared developmental pathophysiology for ARBs with phenotypic manifestation determined by the affected basal ganglia cortico-striatal loop(s). This idea is further corroborated by the high level of overlap in known or suspected biomarkers of disorders associated with stereotypy and early-life onset, compared with disorders lacking stereotypies or with late-life onset. Given the homologous, highly conserved nature of basal ganglia circuitry across species (6,29), it is likely that the identified proteomic biomarkers of plasma GSH level and stereotypy in mice would also be predictive of stereotypies and REDOX imbalance in humans. Further investigation into the causal relationship between REDOX imbalance and stereotypy, and translation to other species (e.g. humans), is required to confirm the generality of such results.

The current study had several limitations that warrant comment. First, we used animals retired from other studies rather than specifically breeding animals for our cohorts. While there are arguably disadvantages to this approach, this enrollment method did confer three important benefits: 1) we were able to obtain a highly heterogeneous experimental population, which increases generalizability and reproducibility of results (34,35); 2) we were able to obtain a diverse spread in both stereotypy severity and age without specifically breeding animals and then inducing stereotypies (consistent with the 3Rs) (40); and 3) this design more closely mimics human studies where individual variability is embraced, and unwanted “noise” is dealt with statistically rather than experimentally (35). A second limitation of this study is that due to the opportunistic use of banked blood samples for the CD1 replication cohort, this group of animals consisted of only young animals, which all happened to be females. Additionally, we were unable to quantify plasma GSH level in these animals (due to the nature of how the banked blood samples had been collected) and therefore cannot confirm the relationship between proteomic expression and plasma GSH level in this second cohort. Finally, although the Olink platform is compatible with both heparin and EDTA anticoagulants, it is important to note that different anticoagulants were used for sample collection between the two cohorts—heparin was used for C57BL/6 mice and EDTA was used for CD1 mice. Despite this, there was still a significant and robust link between stereotypy and proteomic expression in both cohorts.

These novel findings suggest multiple directions for future research. These include: 1) determining whether plasma GSH level and/or the identified proteomic biomarkers of plasma GSH level and severity of stereotypy are present in other species (e.g., in nonhuman primates and human clinical populations); 2) investigating whether plasma GSH level and/or the plasma proteomic biomarker signatures can be used to screen mice for onset of stereotypies 3) determining whether antioxidant therapies, which are effective in treatment of other ARBs (26–28) are also effective in treating and preventing development of stereotypies in mice.

In conclusion, here we establish the relationship between plasma GSH level and stereotypy severity in C57BL/6 mice. We then used a state-of-the art proteomics approach to identify a broader biomarker profile of dysfunction. We found expression levels of nine proteins to correlate with plasma GSH level and expression of 15 proteins to correlate with stereotypy severity. Further supporting REDOX imbalance in the developmental pathophysiology of stereotypies, a large portion of the identified proteins have functional roles in REDOX physiology and are either known, or suspected, biomarkers of neurodevelopmental and neuropsychiatric disorders characterized by presence of stereotypy. Furthermore, given the highly conserved nature of the proteins assessed by the Olink T96 Mouse Exploratory Panel, these results have promising translational potential for clinical populations. Along with the established relationship between REDOX imbalance and compulsive behaviors (26–28), these data suggest REDOX imbalance as a shared pathophysiology across ARBs and highlight promising novel targets for early detection and intervention.

## Methods

### Ethics Declarations

All experimental procedures were approved by and conducted in accordance with the Stanford University IACUC. Stanford is an AAALAC accredited institution. Additionally, all methods are reported in accordance with ARRIVE guidelines for the reporting of animal experiments.

### Animals and Housing

#### C57BL/6 mice

To ensure a spread of well-developed stereotypies, we used 20 mice (N=10 females; N=10 males) on a C57Bl/6 background ranging in age from 140 to 225 days old that were slated for euthanasia. We singly housed subjects in standard individually ventilated cages (Innovive, San Diego, CA) with 30g of Alphadri bedding and 8g of ALPHA-twist nesting enrichment (Shepard Specialty Papers, Watertown, TN) for the duration of the study. Animals were maintained on a 12hr/12hr light/dark cycle (lights on at 13:15) in a climate-controlled room (72 ± 2°F) with food (Teklad 2018 mouse diet) and deionized water available *ad libitum*. Cages were changed weekly.

#### CD1 mice

This cohort consisted of 28 CD1 retired sentinel mice (all females) ranging in age from 119 to 156 days old that were slated for euthanasia. Animals were housed under the same conditions as described for the C57BL/6 mice.

### Procedures

#### Behavioral Recording and Scoring Stereotypy Severity

Following a seven-day acclimation period, we performed cage changes and unobtrusively recorded home-cage behavior of subjects using a camera system (LaView™ security; City of Industry, CA). Using a previously validated mouse ethogram [See supplementary table S5 online], a trained observer scored 24-hours of home cage behavior for stereotypy severity. Video was broken into 15 minutes bins; the first 5 min of each bin was scored. Each behavior on the ethogram was given a score of either one (1), present within the 5-min observation interval, or zero (0), not present within the observation interval (41). We then summed counts of stereotypies over the 24-hour period to produce a final “total stereotypies” count. This final count of “total stereotypies” provided a measure of stereotypy severity and was hence used in all analyses. To ensure coding was unbiased, the observer was blinded to all measures of glutathione and the proteomic profile of subjects.

### Sample Collection and Processing

Subjects were deeply anesthetized using isoflurane and whole blood collected via cardiac puncture into a syringe; animals were secondarily euthanized via cervical dislocation. Whole blood was transferred into an anticoagulant coated microtainer and stored on ice for transport to the laboratory where it was centrifuged at 4°C for 5 min at 15000 rpm to obtain plasma. For C57BL/6 mice, whole blood was transferred into heparin coated microtainers (BD Microtainer item 13-680-62). The resulting plasma was split into two aliquots: one aliquot was immediately flash frozen on dry ice and stored at -80°C for future analysis with the Olink proteomics panel; the other aliquot was prepared and stored for plasma GSH quantification. The plasma in this aliquot was mixed with an equal volume of 5% aqueous 5-sulfosalicylic Acid (SSA; Sigma Aldrich Item s2130) and incubated for 10 minutes at 4°C, and then centrifuged again at 4°C for 10 min at 14000 rpm. The resulting 2.5% SSA plasma supernatant was collected, flash frozen, and stored at -80°C for future GSH analysis. One subject was excluded from analysis due to hemolysis of sample (n = 19). For CD1 mice, we opportunistically used banked plasma samples where whole blood had been transferred into EDTA coated microtainers (BD microtainer item number 365972-1) prior to centrifugation. The resulting EDTA plasma was flash frozen on dry ice and stored at -80°C.

### GSH Quantification

Plasma level of total GSH, free GSH, and GSSG were assessed using the DetectX glutathione detection kit (#K006, Arbor Assay, Ann Arbor, MI, USA). Following manufacturer instructions, 2.5% SSA plasma was further diluted 1:2.5 with assay buffer to produce a final dilution of 1:5 before assaying. Given prior experience with this kit (26,42), we placed the 96-well plate on an orbital shaker during incubation periods to ensure proper mixing of samples and reagents. Three of the samples had detectable values for total GSH but were below the limit of detection for free GSH. Therefore, we corrected for the missing values by adding the lowest read value of total GSH and its respective value of free GSH to all samples. Based on work conducted in our laboratory, total plasma GSH levels are more reliable than free GSH, GSSG, or the relative ratio (42). As such, total plasma GSH was used in all analyses.

### Olink Proteomic Expression

Plasma samples were transported to the Human Immune Monitoring Center at Stanford University for analysis using the Olink Mouse Exploratory Panel (43). This panel uses a proximity extension assay to simultaneously quantify expression level of 92 proteins (quantified as a normalized expression value (NPX) on a log_2_ scale). Raw data showed evidence of concentration effects. We therefore normalized NPX within subject by averaging the values for all proteins and subtracting this from each individual protein. The within-subject normalized NPX values were used for further analysis.

### Proteomic Ontology

To classify protein function, we conducted individual literature searches for each protein that correlated with GSH or stereotypy severity and passed BH correction for q < 20 in at least one analysis. Proteins were included within functional categories (dopamine, REDOX, and/or neurodevelopmental or neuropsychiatric condition—ASD, schizophrenia, ADHD) if the literature search resulted in at least one publication suggesting linkage between the protein and functional category, and/or the protein was included in databases associated with the disorders of interest (e.g. GeneCards, SFARI).

### Statistical Methods

Initial analyses were conducted using JMP16 Pro for Windows using LS-GLM or REML mixed models, with transformations conducted as necessary to meet assumptions of linear models (homogeneity of variance, normality of error, linearity) (44). Non-significant interactions were removed when doing so improved the AICc significantly, or when a model had high R^2^ but no significant effects (indicating a colinear model). Non-significant interactions were retained when doing so improved the AICc of the model, or when they were conceptually necessary to test a hypothesis or provide a positive control. Further *post-hoc* tests (as described below) were performed in equivalent models using SAS 9.4 for windows.

### GSH and Stereotypy Severity in C57BL/6 Mice

We used a LS-GLM blocked by age, sex, and strain to examine the relationship between plasma GSH level and stereotypy severity. We included both the main effect of GSH level and the interaction between plasma GSH level and age. To meet GLM assumptions, plasma GSH level required a log transformation, and stereotypies required a square root transformation. Initially analyses were conducted using age in days (i.e., as a continuous variable). Upon further examination, we noted that animals grouped distinctly into two age categories. We therefore performed the analyses again using age as a dichotomous variable and found this to be a slightly better model (AICc difference of 2.5). Therefore, we proceeded with age as a dichotomous variable in the analyses.

### Proteomics and GSH in C57BL/6 Mice

We used an REML repeated measures mixed model, with mouse nested within age group as a random effect. This approach controls for subject, including all related non-specified variables (e.g., sex or cage location). Age group, protein, and plasma GSH level were included as fixed effects. We additionally included interactions between age group and protein, age group and plasma GSH level, protein and plasma GSH level, and the three-way interaction between age group, protein, and plasma GSH level. This approach is equivalent to a compositional analysis without the constraint of scaling protein expression to a percentage. Thus, “protein” describes the average expression profile across all subjects, while interactions with protein test how this profile changes with age, GSH level, and their interaction. This approach maximizes power (by accounting for between-subject effects) and minimizes the chance of false discovery (because each question is tested by a single family-level gate-keeping p-value). Finding a significant three-way interaction, we then conducted post-hoc analysis in SAS to examine relationships of individual proteins and GSH level within the two age groups. We used a BH correction to prevent false discovery.

### Proteomics and Severity of Stereotypy in C57BL/6 Mice

Maintaining consistency across analyses, we used the same REML mixed model approach controlled for all individual differences (age was nested within individual as a random effect), to examine the main effects of age group, protein, and stereotypy severity. We also included interactions between age and protein, age group and severity stereotypies, protein and severity stereotypies, and the three-way interaction between age group, protein, and stereotypy severity. This three-way interaction was not significant; however, removal of the interaction term resulted in an increased AICc, and, therefore, was retained in the model. Additionally, retaining this interaction term ensured consistency with the plasma GSH dataset and controlled for any effects due to age group. Finding a significant interaction between stereotypy severity and protein expression, we performed a BH correction to prevent false discovery. For purposes of this analysis, we performed a log transformation on stereotypy severity.

### Proteomic Validation in CD1 Mice

We used a REML mixed model controlled for individual differences, to examine the relationship between main effects of protein and stereotypy severity, and the interaction between protein and stereotypy severity. As all CD1 mice fell within the age range of the young group of mice in the first cohort, age was not included as a main effect. Finding a significant interaction between stereotypy severity and protein expression, we examined correlations between expression of individual proteins and stereotypies and performed a BH correction to prevent false discovery.

For the purpose of this analysis, we performed a log transformation on stereotypies.

## Data Availability

Data and SAS code used for analysis are available in supplementary information (see supplementary information S6 online).

## Acknowledgements

The authors would like to thank the Stanford Veterinary Service Center for assisting with the care and wellbeing of animals used in this study and for assisting with blood collection. Finally, we would like to acknowledge the Stanford University Discovery Innovation Fund for providing financial support for this research.

## Author Contributions

KMC, KJP, and JPG designed the study; KMC, KJB, BESB, KAB, and JNB performed research; JTG and RM provided research support; KMC, KJB, and JPG analyzed data; KMC, KJP, and JPG wrote the manuscript; all authors edited and approved the final manuscript.

